# Seasonal variation in D2/3 dopamine receptor availability in the human brain

**DOI:** 10.1101/2023.12.20.572517

**Authors:** Lihua Sun, Tuulia Malén, Jouni Tuisku, Valtteri Kaasinen, Jarmo A. Hietala, Juha Rinne, Pirjo Nuutila, Lauri Nummenmaa

**Affiliations:** Turku PET Centre, University of Turku, Turku, Finland; Turku PET Centre, Turku University Hospital, Turku, Finland; Huashan Hospital, Fudan University, Shanghai, China; Clinical Neurosciences, University of Turku, Turku, Finland; Neurocenter, Turku University Hospital, Turku, Finland; Department of Psychiatry, University of Turku and Turku University Hospital, Turku, Finland; Department of Endocrinology, Turku University Hospital, Finland; Department of Psychology, University of Turku, Turku, Finland

**Keywords:** Dopamine D2 receptor, seasonality, striatum, caudate, positron emission tomography, reward

## Abstract

Brain functional and physiological plasticity is essential to combat dynamic environmental challenges. The rhythmic *in vivo* dopamine signaling pathway, which regulates emotion, reward and learning, shows seasonal patterns with higher capacity of dopamine synthesis and lower number of dopamine transporters during dark seasons. However, seasonal variation of the dopamine receptor signaling remains to be characterized. Here, based on historical database of healthy human brain [^11^C]raclopride PET scans (n = 291, 224 males and 67 females), we investigated the seasonal patterns of D2/3 dopamine receptor signaling. We found that daylength at the time of scanning was negatively correlated with availability of this type of receptors in the striatum. Likely, seasonally varying D2/3 receptor signaling also underlies the seasonality of mood, feeding, and motivational processes.

**Significance of the study:** Brain *in vivo* neurotransmitter signaling demonstrates seasonal patterns. The dopamine D2/3 receptor, with both pre- and postsynaptic expressions, has been targeted by major antipsychotic and antiparkinsonian pharmaceuticals. The current study, based on a large dataset of healthy brain PET images quantifying these receptors, shows that dark seasons are associated with increased receptor availability in the striatum. Considering the previous findings of increased dopamine synthesis and lowered number of transporters, findings may suggest elevated presynaptic control of dopamine release and increased dopamine receptor sensitivity during dark seasons. The rhythmic D2/3 receptor availability may be a mechanism underlying seasonality in mood, feeding, and motivational processes.

## Introduction

Brain functional and physiological plasticity is essential to combat dynamic environmental challenges, for survival and the wellbeing. This includes the potential seasonal patterns of dopamine signaling. For instance, *in vivo* imaging data in both healthy subjects (Eisenberg et al., 2010) and patient data (Kaasinen et al., 2012) show increased of striatal dopamine synthesis in fall and winter. Dopamine transporter binding in the left caudate is found lower during dark seasons (Booij et al., 2023). Also, preclinical studies suggest that longer photoperiod stimulates nucleus accumbens dopamine release in female mice (Jameson et al., 2023). Dopamine signaling plays an important role in seasonal breeding of animals (Gerlach and Aurich, 2000) and, via interaction with the melatonin signaling, affects circadian rhythms (Cahill and Besharse, 1991; El Halawani et al., 2009). Further, dopamine regulates emotion, reward, and learning that demonstrate seasonal patterns, with learning ability, attention and positive emotions all at low levels during winter months (Golder and Macy, 2011; Meyer et al., 2016; Mooldijk et al., 2022). Human feeding behaviors, where the brain dopamine signaling also plays a crucial role (Small et al., 2003), similarly demonstrate seasonal patterns with increased caloric intake of fats often found in fall or winter (Shahar et al., 1999). Detailed knowledge on how dopamine signaling adapts to seasonal rhythms is not only essential for understanding normal molecular brain plasticity, but also crucial in understanding psychiatric conditions with seasonally varying onset and severity, such as the seasonal affective disorders.

Brain dopamine signaling is primarily relayed through the D1 family and D2 family receptors (D2Rs), where the D2Rs are found with both post- and pre-synaptic expression. This indicates an additional role of D2Rs in receptor-mediated feedback control in dopamine signaling. D2Rs are involved in both presynaptic dopamine release (Liu and Kaeser, 2019) and in regulation of dopamine synthesis (Sulzer et al., 2016). For instance, D2R agonist inhibits tyrosine hydroxylase (TH), where TH action on L-DOPA is a rate-limiting step for dopamine synthesis (Pothos et al., 1998). Also, cAMP is known to induce expression of TH, and D2Rs inhibit the cAMP signaling (Chen et al., 2008). Further, D2Rs may possess complementary interaction with dopamine transporter, with evidence showing that D2Rs enhance the velocity of dopamine synaptic reuptake (Schmitz et al., 2002; Wu et al., 2002). Therefore, rhythmic D2R functions potentially direct all steps of dopamine signaling including its presynaptic synthesis and release, and extracellular clearance. Yet, potential seasonal patterns of D2R signaling remains to be characterized.

Here we analyzed large historical dataset of healthy subjects’ brain PET scans (n = 291) measuring D2Rs (i.e., primarily the D2/3 receptors) with antagonist radioligand [^11^C]raclopride. We focused on striatal and thalamic areas where this receptor is highly expressed and most reliably measured with the used radioligand (Hirvonen et al., 2003; Freiburghaus et al., 2021). Daylength was used as a regressor to predict the regional D2R availability, and hemispheric brain data were analyzed separately. Prior findings show that dark seasons are associated with enhanced dopamine synthetic ability (Eisenberg et al., 2010; Kaasinen et al., 2012) and also reduced amount of dopamine transporter (Booij et al., 2023). This may indicate reduced amount of extracellular dopamine, despite of increased synthesis, and enhanced presynaptic control of dopamine release. Therefore, considering the autoreceptor role of D2Rs in a feedback control, we hypothesized short daylength to be associated with increased striatal D2R availability.

## Methods

### Data

The data were 291 baseline [^11^C]raclopride scans from healthy control subjects (224 males and 67 females; age 19-81 years) collected at Turku PET Centre between 2004 and 2018. Data distribution to local daylength and age are illustrated in **Fig. 1**. For each subject, daylength was calculated as the daytime plus civil twilight on the day when the PET image was acquired, based on geographic location of the Turku PET Center (Turku, Finland; latitude = 60.4518; longitude = −22.2666), as previously (Sun et al., 2021).

**Figure 1.**
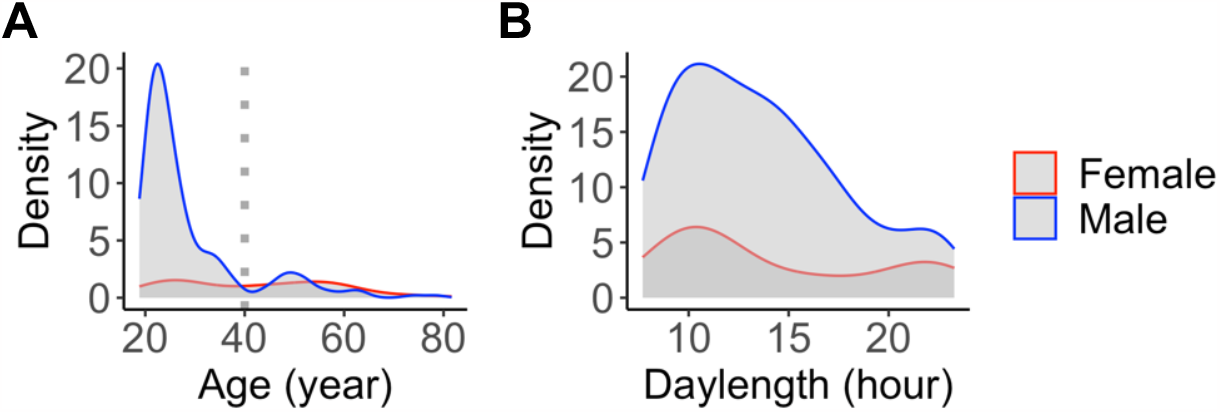
Sex-specific age and local daylength distributions (densities) at the time of scanning. Dotted line shows the age cutoff (40 years) for the primary sample.

### PET data acquisition and image processing

Antagonist radioligand [^11^C]raclopride binds to D2Rs, allowing reliable quantification of striatal and thalamic D2R availability (Farde et al., 1985). In this study, we included the following regions of interest (ROIs) as delineated based on the AAL atlas (Rolls et al., 2015): nucleus accumbens, caudate nucleus (caudate), putamen, as well as thalamus, which were separately analyzed for left and right hemispheres.

Preprocessing was done using the Magia toolbox (Karjalainen et al., 2020). Tracer binding was quantified using the outcome measure binding potential (*BP*_*ND*_), which is the ratio of specific binding to non-displaceable binding in tissue (Innis, 2007). *BP*_*ND*_ was estimated using a simplified reference tissue model (SRTM) (Lammertsma and Hume, 1996) with cerebellar gray matter as the reference region. Acquisition length was harmonized by including first 52 minutes from each scan (Hirvonen et al., 2003), independently of their scan duration.

### Statistical analysis

Data were analyzed using linear mixed effects regression, via the R statistical software (version 4.3.0) and the lme4 package. Because ageing influences D2R availability (Sowell et al., 2003; Collier et al., 2017; Malén et al., 2022), primary analysis was based on subjects younger than 40 years old (primary analysis, n = 227). Findings based on the whole sample (n = 291) are presented in the supplementary data.

D2R *BP*_*ND*_ was modelled separately for each region of interest using fixed factors including daylength at scanning, age and sex; scanner types were used as the random intercept. D2R *BP*_*ND*_ were log-transformed, while daylength and age were standardized in the statistical models. Because BMI is only weakly associated with *BP*_*ND*_ (Malén et al., 2022) and a large portion of the subjects lacked BMI data, it was excluded from the statistical models.

### Voxel-level analysis

The data were analyzed at the voxel level using SPM12 (Wellcome Trust Center for Imaging, London, UK, http://www.fil.ion.ucl.ac.uk/spm). The normalized *BP*_*ND*_ images were entered into general linear models, where they were predicted with daylength. Age, sex and scanner types were entered into the models as nuisance covariates. Because [^11^C]raclopride binds selectively only in striatum and thalamus, the analysis restricted to the high-binding sites by creating a single mask covering these regions (caudate, putamen, nucleus accumbens and thalamus). Statistical threshold was set at *p < 0*.*05*, FDR-corrected at cluster level.

## Results

### Subjects with age under 40 years old

The primary sample (n = 227) included 195 males and 32 females. The mean age was 25.6 y (SD = 4.87 y, range = 18.82 – 39.47 y) and average daylength exposure was 13.61 h (SD = 4.26 h, range = 7.7 – 23.25 h). More detailed information of the primary sample is found in *supplementary Table S1*.

Data revealed that daylength was linearly and negatively associated with regional D2R availability (*BP*_*ND*_) in most regions (**Table 1, Fig. 2 & Fig. 3)**. Age was negatively associated with D2R *BP*_*ND*_, and male subjects had lower D2R *BP*_*ND*_ (**Fig. 2**) than females. Information of the full sample and the corresponding findings are found in *supplementary Table S2 & S3*.

**Table 1.** Effect of daylength on regional D2R availability in the brain (uncorrected for multiple comparison).

| Hemisphere | Region | Beta | 95% CI | t | p |
| --- | --- | --- | --- | --- | --- |
| Left | Caudate | -0.028 | -0.042, -0.014 | -3.86 | 0.00015*** |
| Right | Caudate | -0.018 | -0.032, -0.0033 | -2.41 | 0.017* |
| Left | Putamen | -0.013 | -0.025, -0.001 | -2.11 | 0.036* |
| Right | Putamen | -0.012 | -0.024, -0.000006 | -1.95 | 0.053* |
| Left | NACC | -0.019 | -0.036, -0.0024 | -2.23 | 0.027* |
| Right | NACC | -0.016 | -0.033, 0.0015 | -1.78 | 0.077 |
| Left | Thalamus | -0.016 | -0.039, 0.0056 | -1.46 | 0.15 |
| Right | Thalamus | -0.0065 | -0.028, 0.015 | -0.60 | 0.55 |

**Figure 2.**
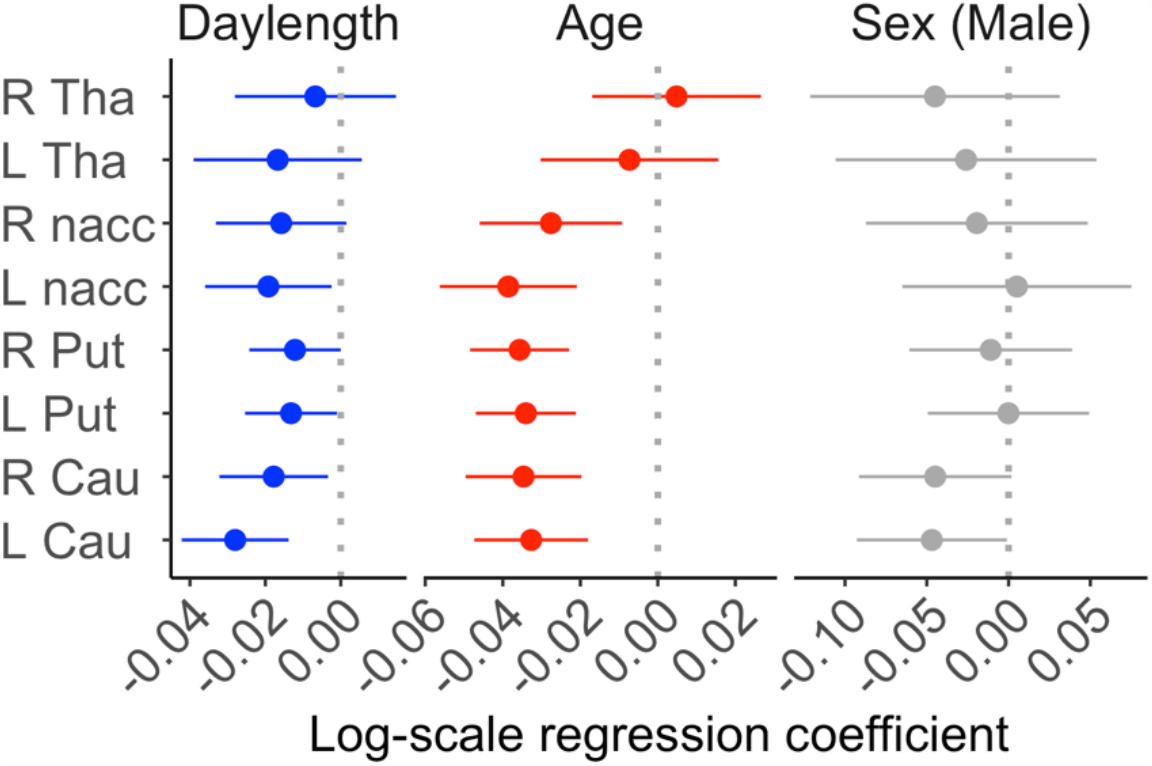
Effect sizes (i.e., point estimate and the 95% confidence interval) of daylength, age and sex (male) on D2R BP_ND_ in different brain regions. L = Left, R = Right, Cau = Caudate, Put = Putamen, nacc = Nucleus accumbens, Tha = Thalamus.

**Figure 3.**
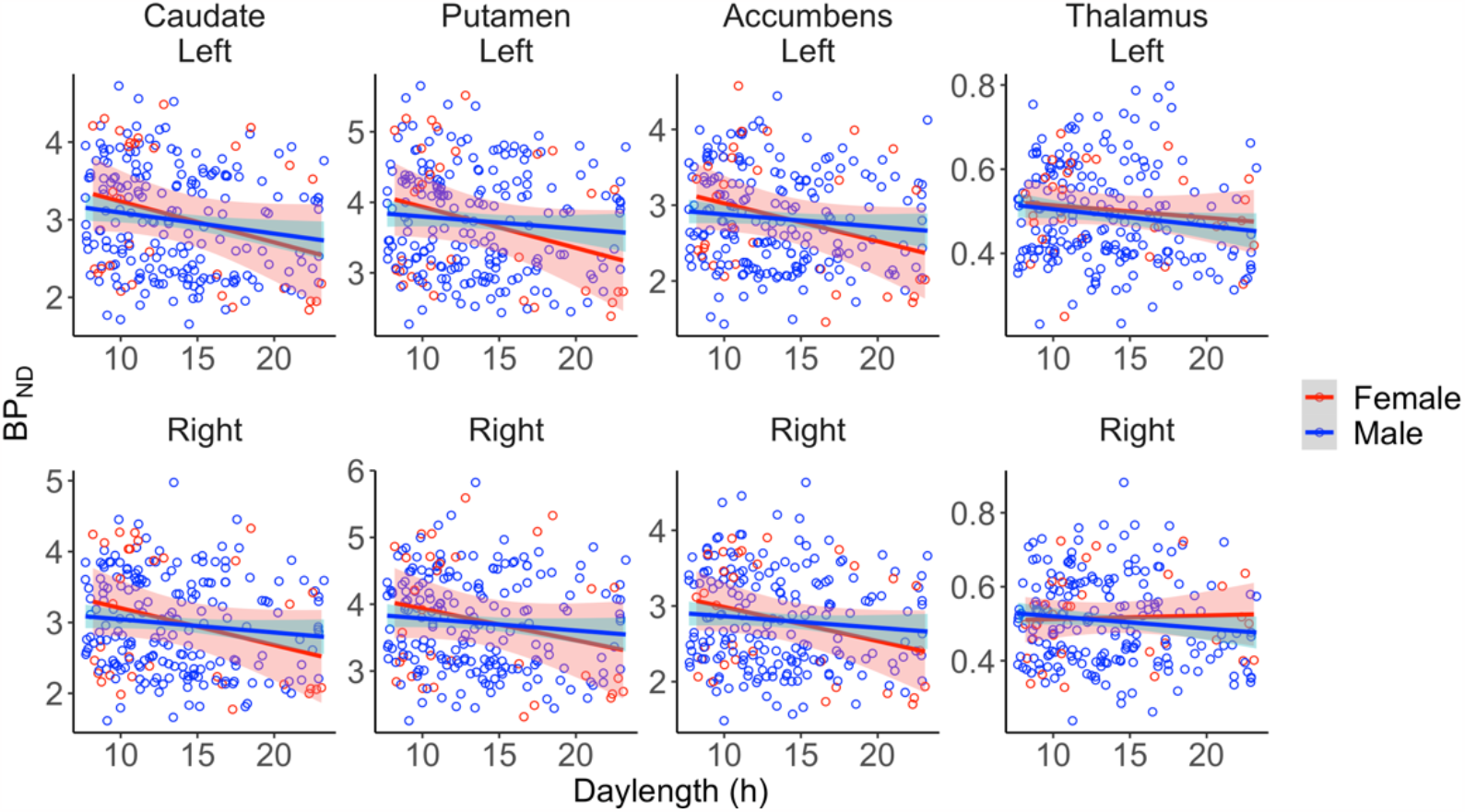
Association between regional D2R BP_ND_ and daylength in each ROI, separately for males and females. Red and blue lines show Least Squares regression lines, and their 95% confidence intervals are shaded.

We compared the effect size (i.e., 95% CI) between daylength, age and sex, **Fig 2**. In the primary sample, every 4.26 h (i.e., one standard deviation) increase of daylength was associated with a mean 2.8% drop of D2R *BP*_*ND*_ in the left caudate. In the same region, every 4.87 h (i.e., one standard deviation) increase of age was associated with a mean 3.3% drop of D2R *BP*_*ND*_. Corresponding results based on the full sample are found in *Supplementary Figure S1*.

### Voxel-level analysis of the primary sample

We next run a complementary voxel-level analysis for the high-binding regions (striatum and thalamus). Similar as in the ROI analysis, Daylength was found to be a significant predictor for striatal D2R *BP*_*ND*_, **Fig 4**.

**Figure 4.**
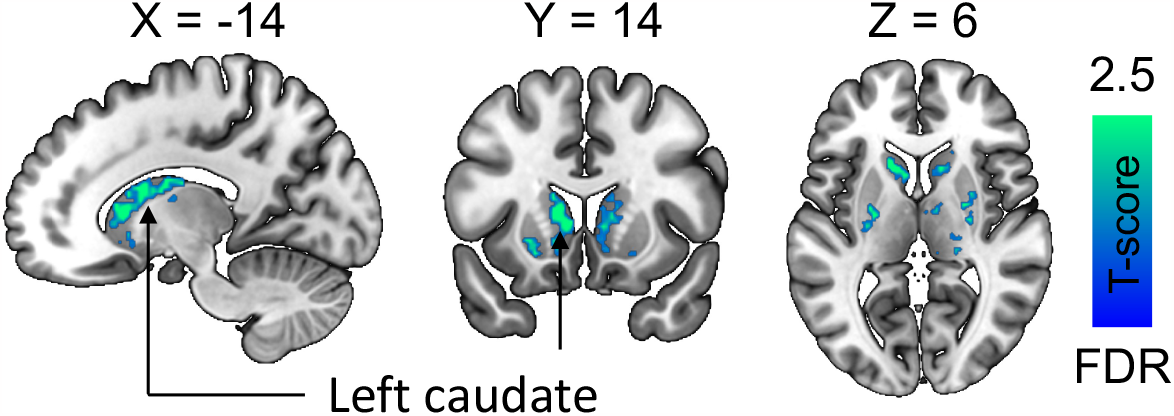
Daylength was as significant predictor for D2R BP_ND_ in the striatum. Data are thresholded at P < 0.05 with false discovery rate (FDR) cluster-level correction.

### Effect of daylength across age cohorts

Considering the impact of ageing on dopamine signaling (Sowell et al., 2003; Collier et al., 2017; Malén et al., 2022), there is always a tradeoff between sample size and the allowed maximum age. In addition to a primary sample analysis, we also investigated how the effect of daylength survived a stepwise minimization of sample in accord to the allowed maximum age.

We studied the effect of daylength in different groups where the whole sample was divided into 7 groups in accord to the allowed maximum age, **Table 2**. Our data showed that the effect of Daylength of D2R *BP*_*ND*_ retained across different groups, especially in the left caudate (**Fig. 5** and *Supplementary Figure S2*). The finding goes in line with previous findings that highlight the importance of the left caudate region, regarding both seasonal variation of dopamine transporter signaling (Booij et al., 2023) and dopamine-relevant etiology in seasonal affective disorders (Neumeister et al., 2001).

**Table 2.**
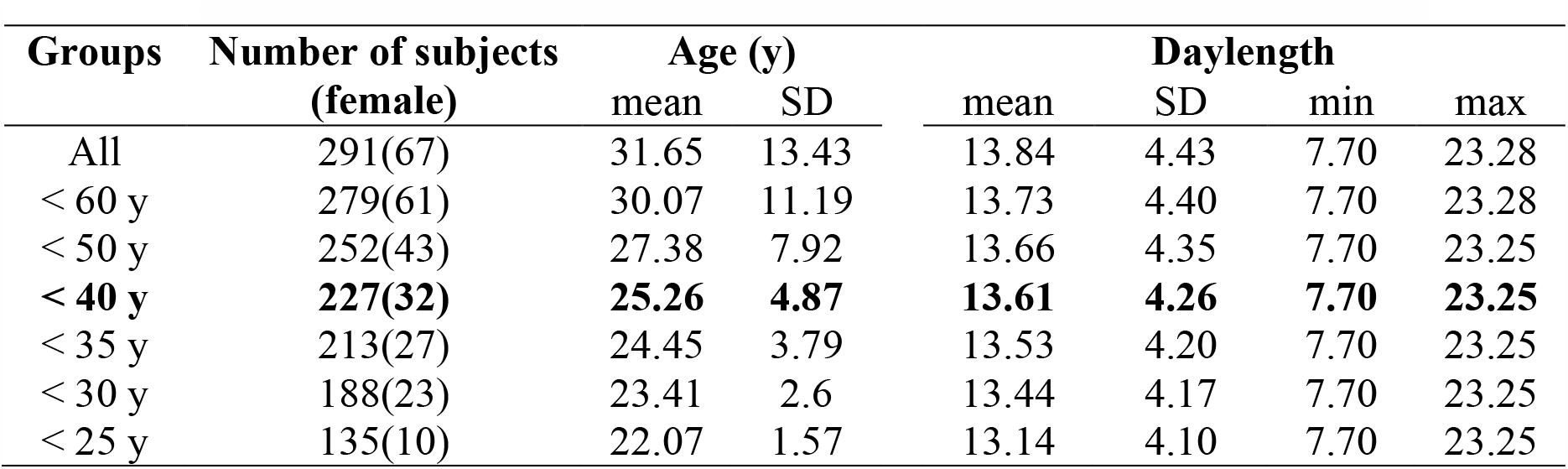
Daylength and age information across groups by age.

| Groups | Number of subjects<br>(female) | Age (y) |  | Daylength |  |  |  |
| --- | --- | --- | --- | --- | --- | --- | --- |
|  |  | mean | SD | mean | SD | min | max |
| All | 291(67) | 31.65 | 13.43 | 13.84 | 4.43 | 7.70 | 23.28 |
| < 60 y | 279(61) | 30.07 | 11.19 | 13.73 | 4.40 | 7.70 | 23.28 |
| < 50 y | 252(43) | 27.38 | 7.92 | 13.66 | 4.35 | 7.70 | 23.25 |
| **< 40 y** | **227(32)** | **25.26** | **4.87** | **13.61** | **4.26** | **7.70** | **23.25** |
| < 35 y | 213(27) | 24.45 | 3.79 | 13.53 | 4.20 | 7.70 | 23.25 |
| < 30 y | 188(23) | 23.41 | 2.6 | 13.44 | 4.17 | 7.70 | 23.25 |
| < 25 y | 135(10) | 22.07 | 1.57 | 13.14 | 4.10 | 7.70 | 23.25 |

**Figure 5.**
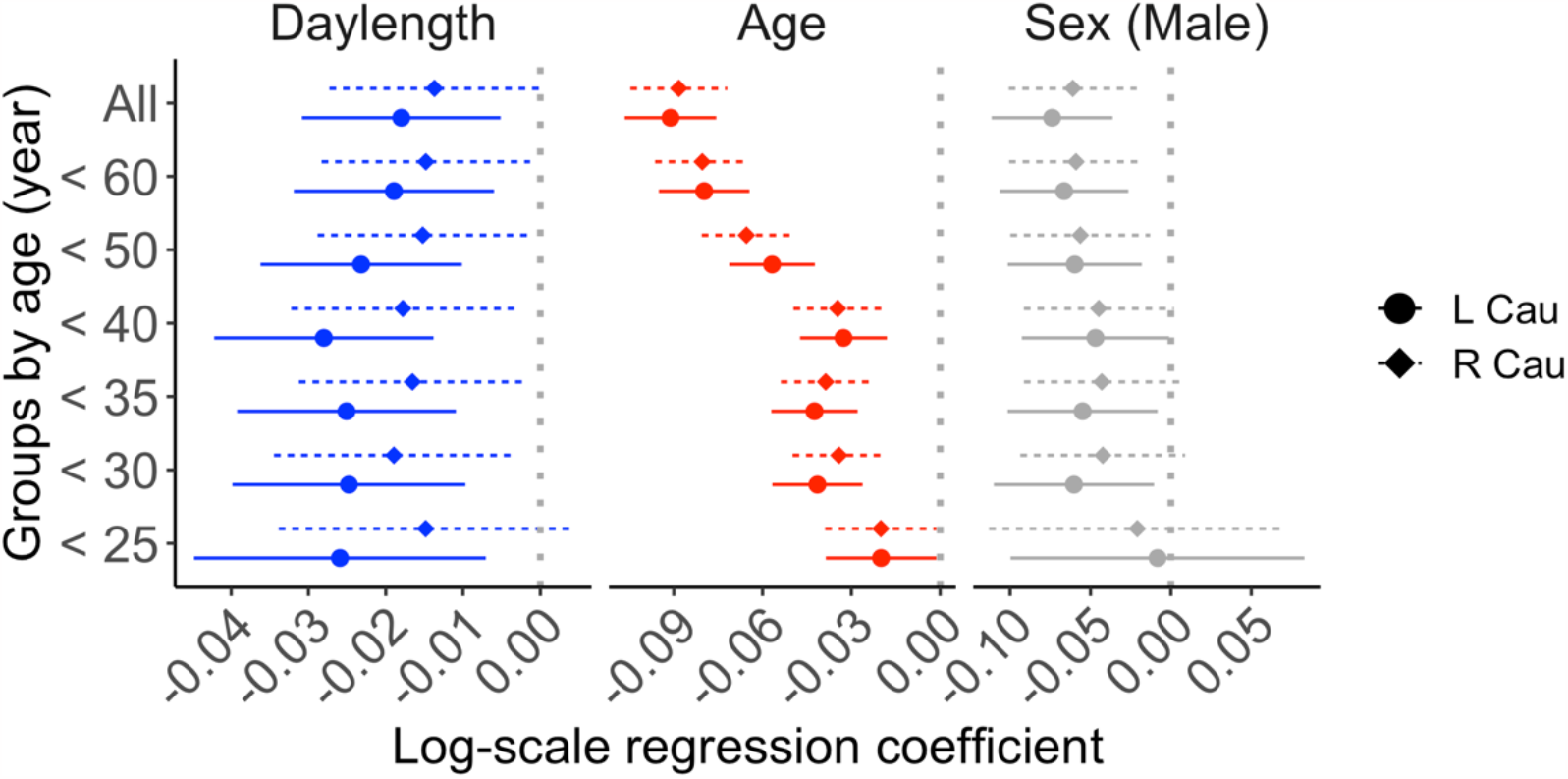
Effect sizes for daylength, age and sex (male) on D2R BP_ND_ in the left and right caudate in different groups defined by maximum age. Plots of other ROIs are found in the supplementary Figure S2.

## Discussion

Our main finding was that daylength modulated *in vivo* brain D2R availability in healthy humans. Specifically, striatal D2R availability was elevated in autumn-winter time when the days are short and lowered during the spring-summer time when the days are long. This finding is based on, to our knowledge, the largest database of healthy human D2R PET images. The pattern of seasonal change was the most consistent in the left caudate, aligning with previous studies showing that dark seasons are associated with lowered amount of dopamine transporter in the left caudate (Booij et al., 2023). Also, patients with seasonal affective disorder (SAD) show lowered dopamine transporter in the left caudate (Neumeister et al., 2001), possibly indicating that dopamine signaling in the left caudate is linked with seasonal onsets of depression symptoms. The effect was also strong: one standard deviation of daylength change (i.e., around 4 h) was associated with equal magnitude of D2R binding changes as 2-5 years of ageing in the left caudate. The effect of daylength on D2R was comparable in the datasets with only young and middle-aged adults as well as in the data with full age range. This suggests that the impact of daylength is consistent across age cohorts. Altogether these data suggest that in future studies of D2R binding, especially in high-latitude regions, the effect of seasonality should be considered as it may confound the primary results.

Seasonality in human brain physiology and particularly neurotransmission remains poorly characterized (Levitan, 2022; Zhang and Volkow, 2023). There is evidence showing that dark seasons are associated with lowered postsynaptic serotonin receptor availability (Spindelegger et al., 2012; Matheson et al., 2015) and most probably increased serotonin transporter binding (Koskela et al., 2008; Praschak-Rieder et al., 2008). Clinical *in vivo* data also indicates a light-sensitive fluctuation in levels of cerebral monoamine oxidase A, an enzyme that degrades neurotransmitters including serotonin and dopamine (Spies et al., 2018). Similarly, a paucity of studies on neuropeptide signaling suggest that the endogenous opioid signaling responds to seasonal rhythms (Sun et al., 2021; Sun et al., 2022). The present results on dopamine receptor signaling further highlight the brain neurotransmission mechanism on seasonal patterns of cognitive and affective functions.

Previous studies of dopamine signaling using ^18^F-DOPA PET have found an increase in presynaptic synthesis of dopamine during autumn and wintertime (Eisenberg et al., 2010; Kaasinen et al., 2012). This complements our study showing increased D2R availability during dark season, signifying that increased D2R availability most probably indicates enhanced presynaptic control of dopamine release. Also, D2Rs are known as autoreceptors that modulate the presynaptic synthesis of dopamine (Pothos et al., 1998; Chen et al., 2008; Sulzer et al., 2016), and increased D2R signaling may support this in encountering the increased dopamine synthetic capability. This is also supported by findings that the dopamine transporter in the left caudate is lower during dark seasons (Booij et al., 2023), as D2Rs may restrict dopamine release and enhance transporter functions (Schmitz et al., 2002; Wu et al., 2002). Besides, the increased availability of D2Rs may also be explained by increased melatonin release during dark seasons, since melatonin is known to inhibit presynaptic dopamine release (Wehr, 1991; Zisapel, 2001).

However, one study using single photon emission computed tomography with [^123^I]iodo-benzamide to measure dopamine D2/D3 receptor availability, shows that shorter rather than longer sunlight exposure is associated with reduced amounts of striatal receptors (Tsai et al., 2011). This study used data from low latitude regions where variation of daylength across seasons is small (i.e., around 4 hours between brightest and darkest days in local region), and the subjects are divided into two groups where unbalanced sex and smoker effects may also complicate the interpretation of the finding. Further, reference tissue modeling of PET data in the study uses the frontal lobe as a reference region and this may also compromise their conclusions.

The striatum is a brain key node for reward responses (Balleine et al., 2007) and energy consumption in the caudate may directly code the feeling of satiation or hunger (Sun et al., 2023). Given the role of striatal dopamine signaling in feeding behavior, enhanced D2R signaling in dark seasons may induce overeating (Shahar et al., 1999). However, few studies focused on the role of dopamine signaling in SADs (Neumeister et al., 2001), and it is challenging to interpret how malfunction in a signal component of this signaling pathway predispose specific symptom such as the overeating character of SAD. Seasonal rhythms profoundly impact mood, with negative affect such as depression, anger, and hostility at lowest rate during the summer (Harmatz et al., 2000), whereas symptoms of SAD peak during the winter months (Lam, 2000). These changes are probably mediated by slow phasic changes in a variety neuroreceptor systems including the dopamine receptor signaling.

## Limitations

[^11^C]raclopride *BP*_*ND*_ in a baseline condition is proportional to D2R density, but the exact contributions of D2R density, receptor affinity, and baseline occupancy by endogenous dopamine cannot be assessed in a single measurement. These components cannot be differentiated in a single scan. The study was based on historical data, where each subject was imaged only once, and quasi-experimental design (i.e., natural changes in daylight) was used as longitudinal multi-scan studies would yield a significant radiation load. The data were sampled from different projects and scanners; potential scanner-related biases were however accounted for in the analyses. Daylength was regarded as a noise-free regressor to index local seasons. Day-to-day variance of sunlight exposure and contribution of other seasonal factors (e.g., atmospheric pressure, temperature), however, were not investigated. Also, the human database was compiled from historical data, where relevant behavioral and self-report measures, such as mood and eating habits, were not systematically collected. Finally, current findings based on human data should be cautiously interpolated into regions with lower latitudes, considering the large magnitude of local photoperiodic variation.

## Conclusions

We conclude that striatal D2R availability is increased during dark seasons. While the exact mechanisms and subsequent impacts of this seasonal pattern of receptor signaling cannot be resolved in this cross-sectional study, our study nevertheless reveals brains’ physiological adaption to seasons at the level of single neurotransmitter system. Considering the important role of striatal dopamine signaling in reward-related behaviors, elevated D2Rs may contribute to the elevated food seeking during dark seasons.

## Acknowledgement

The study is supported by the Academy of Finland (#332225), Sigrid Juselius Stiftelse, Signe och Ane Gyllenberg’s stiftelse, and Turku University Hospital. We thank Päivikki and Sakari Sohlberg Foundation, Finnish Governmental Research Funding for Turku University Hospital and Western Finland collaborative area, and the Finnish Brain Foundation (personal grants to TM).

## Supplementary data

**Supplementary Table S1.** Basic information of the primary sample.

| Scanner<br>Types | Sex | Age |  |  |  | Daylength |  |  |  | N |
| --- | --- | --- | --- | --- | --- | --- | --- | --- | --- | --- |
|  |  | *mean* | *s.d.* | *min* | *max* | *mean* | *s.d.* | *min* | *max* |  |
| S1 | M | 24.62 | 5.11 | 19.24 | 34.68 | 14.8 | 3.08 | 9.12 | 19.66 | 13 |
| S2 | M | 26.8 | 5.12 | 19.43 | 38.75 | 14.34 | 3.89 | 7.73 | 22.5 | 61 |
| S3 | M | 23.58 | 3.25 | 19.22 | 36.17 | 13.23 | 3.99 | 7.71 | 23.25 | 88 |
| S4 | M | 22.9 | 4.07 | 18.87 | 35.58 | 10.76 | 3.3 | 7.79 | 22.89 | 18 |
| S5 | M | 27.46 | 5.39 | 20.55 | 37.03 | 14.9 | 5.47 | 7.7 | 22.9 | 15 |
| S1 | F | 31.06 | 9.03 | 19.47 | 39.47 | 14.33 | 6.17 | 8.3 | 22.76 | 8 |
| S2 | F | 26.28 | 3.36 | 20.1 | 30.33 | 15.19 | 6.16 | 8.53 | 23.08 | 8 |
| S3 | F | 26.79 | 2.49 | 25.07 | 32.69 | 14.79 | 5.07 | 9.93 | 22.51 | 8 |
| S4 | F | 18.82 | - | 18.82 | 18.82 | 11.88 | - | 11.88 | 11.88 | 1 |
| S6 | F | 26.94 | 6.34 | 19.82 | 39.12 | 10.83 | 3.1 | 8.19 | 17.49 | 7 |

N = number of subjects, M = male, F = female, s.d. = standard deviation, min = minimum, max = maximum

**Supplementary Table S2.** Basic information of the full sample.

| Scanner<br>Types | Sex | Age |  |  |  | Daylength |  |  |  | N |
| --- | --- | --- | --- | --- | --- | --- | --- | --- | --- | --- |
|  |  | *mean* | *s.d.* | *min* | *max* | *mean* | *s.d.* | *min* | *max* |  |
| S1 | M | 40.14 | 19.65 | 19.24 | 78.4 | 14.95 | 3.84 | 9.12 | 23.09 | 23 |
| S2 | M | 31.89 | 10.83 | 19.43 | 57.78 | 14.4 | 4.03 | 7.72 | 22.5 | 78 |
| S3 | M | 23.58 | 3.25 | 19.22 | 36.17 | 13.23 | 3.99 | 7.71 | 23.25 | 88 |
| S4 | M | 22.9 | 4.07 | 18.87 | 35.58 | 10.76 | 3.3 | 7.79 | 22.89 | 18 |
| S5 | M | 29.57 | 7.83 | 20.55 | 47.12 | 14.34 | 5.4 | 7.7 | 22.9 | 17 |
| S1 | F | 48.76 | 17.05 | 19.47 | 81.62 | 15.84 | 5.38 | 8.3 | 23.28 | 21 |
| S2 | F | 40.64 | 16.25 | 20.1 | 70.76 | 13.97 | 5.41 | 8.53 | 23.13 | 16 |
| S3 | F | 30.26 | 10.67 | 25.07 | 58.02 | 14.39 | 4.89 | 9.93 | 22.51 | 9 |
| S4 | F | 18.82 | NA | 18.82 | 18.82 | 11.88 | NA | 11.88 | 11.88 | 1 |
| S6 | F | 42.2 | 13.02 | 19.82 | 58.67 | 13.08 | 5.13 | 8.19 | 22.92 | 20 |

N.B., N = number of subjects, M = male, F = female, s.d. = standard deviation, min = minimum, max = maximum

**Supplementary Table S3.**
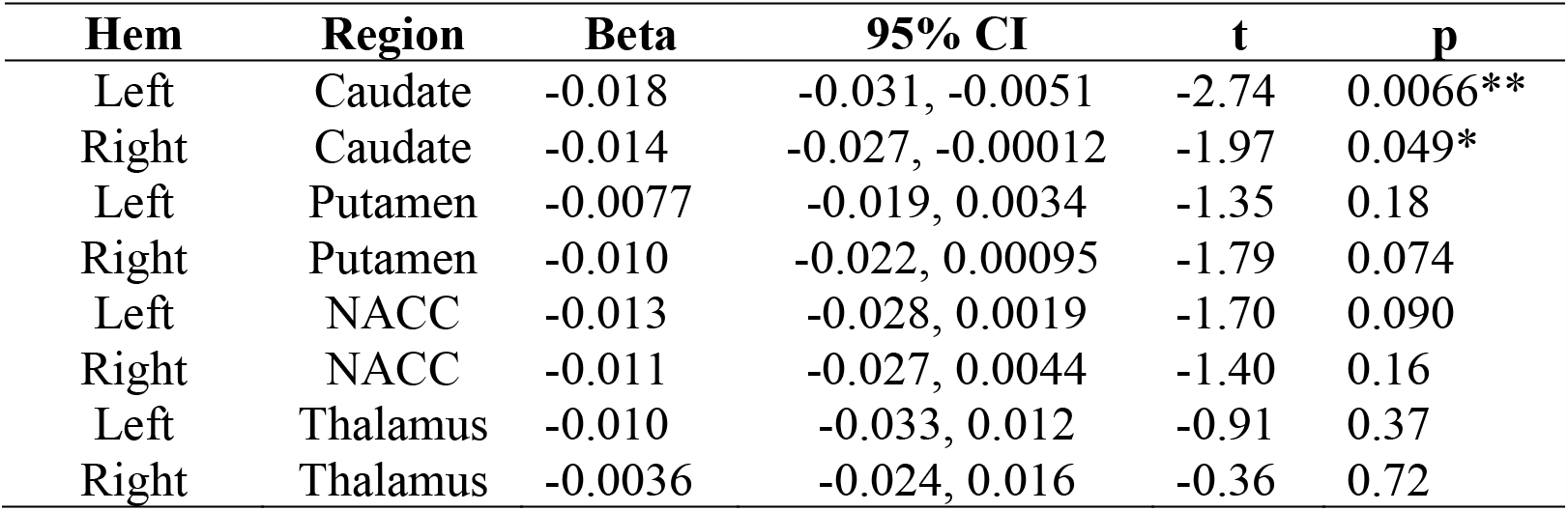
Effect of daylength on regional D2R availability in the full sample (uncorrected for multiple comparison).

| Hem | Region | Beta | 95% CI | t | p |
| --- | --- | --- | --- | --- | --- |
| Left | Caudate | -0.018 | -0.031, -0.0051 | -2.74 | 0.0066** |
| Right | Caudate | -0.014 | -0.027, -0.00012 | -1.97 | 0.049* |
| Left | Putamen | -0.0077 | -0.019, 0.0034 | -1.35 | 0.18 |
| Right | Putamen | -0.010 | -0.022, 0.00095 | -1.79 | 0.074 |
| Left | NACC | -0.013 | -0.028, 0.0019 | -1.70 | 0.090 |
| Right | NACC | -0.011 | -0.027, 0.0044 | -1.40 | 0.16 |
| Left | Thalamus | -0.010 | -0.033, 0.012 | -0.91 | 0.37 |
| Right | Thalamus | -0.0036 | -0.024, 0.016 | -0.36 | 0.72 |

**Supplementary Figure S1.**
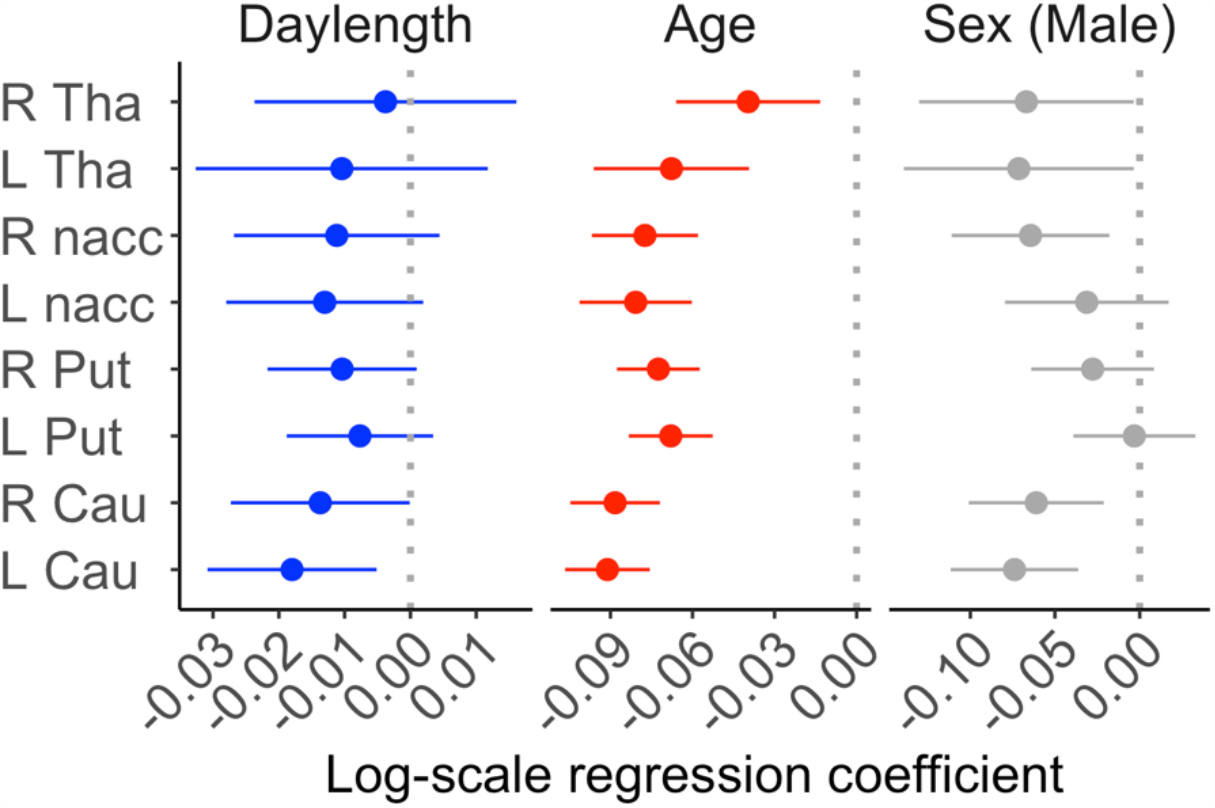
Effect sizes of daylength, age and sex (male) on D2R BP_ND_ in different brain regions in the full sample. L = Left, R = Right, Cau = Caudate, Put = Putamen, nacc = Nucleus accumbens, Tha = Thalamus.

**Supplementary Figure S2.**
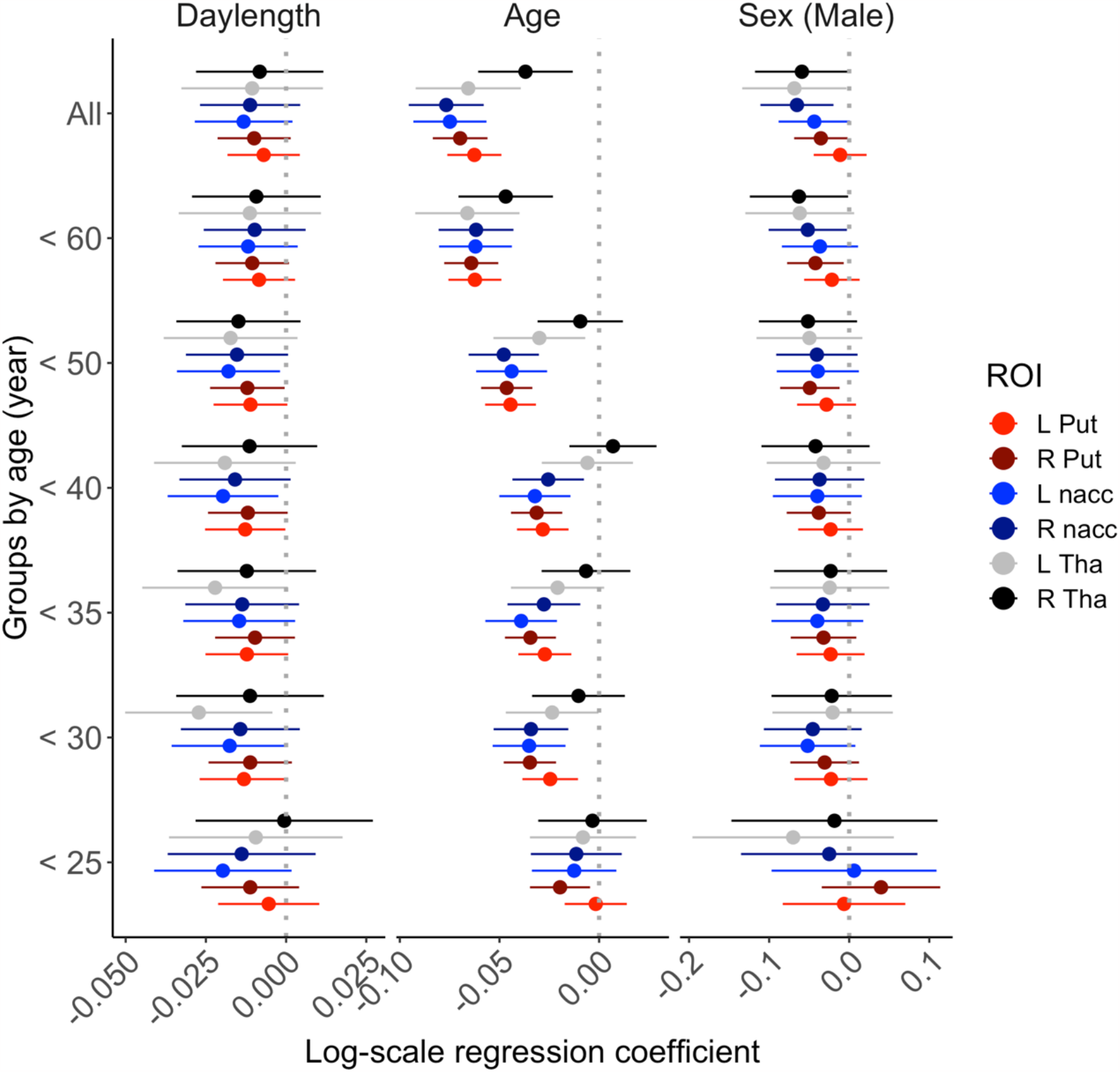
Effect sizes for daylength, age and sex (male) on D2R BP_ND_ in different groups defined by maximum age. ROI = region of interest.

## References

Balleine BW, Delgado MR, Hikosaka O (2007) The role of the dorsal striatum in reward and decision-making. Journal of Neuroscience 27:8161–8165.

Booij J, Tellier SP, Seibyl J, Vriend C (2023) Dopamine Transporter Availability in Early Parkinson’s Disease is Dependent on Sunlight Exposure. Movement disorders.

Cahill GM, Besharse JC (1991) Resetting the circadian clock in cultured Xenopus eyecups: regulation of retinal melatonin rhythms by light and D2 dopamine receptors. Journal of Neuroscience 11:2959–2971.

Chen X, Xu L, Radcliffe P, Sun B, Tank AW (2008) Activation of tyrosine hydroxylase mRNA translation by cAMP in midbrain dopaminergic neurons. Molecular pharmacology 73:1816–1828.

Collier TJ, Kanaan NM, Kordower JH (2017) Aging and Parkinson’s disease: different sides of the same coin? Movement Disorders 32:983–990.

Eisenberg DP, Kohn PD, Baller EB, Bronstein JA, Masdeu JC, Berman KF (2010) Seasonal effects on human striatal presynaptic dopamine synthesis. Journal of Neuroscience 30:14691–14694.

El Halawani ME, Kang SW, Leclerc B, Kosonsiriluk S, Chaiseha Y (2009) Dopamine–melatonin neurons in the avian hypothalamus and their role as photoperiodic clocks. General and comparative endocrinology 163:123–127.

Farde L, Ehrin E, Eriksson L, Greitz T, Hall H, Hedström C, Litton J-E, Sedvall G (1985) Substituted benzamides as ligands for visualization of dopamine receptor binding in the human brain by positron emission tomography. Proceedings of the National Academy of Sciences 82:3863–3867.

Freiburghaus T, Svensson JE, Matheson GJ, Plavén-Sigray P, Lundberg J, Farde L, Cervenka S (2021) Low convergent validity of [11C] raclopride binding in extrastriatal brain regions: A PET study of within-subject correlations with [11C] FLB 457. NeuroImage 226:117523.

Gerlach T, Aurich JE (2000) Regulation of seasonal reproductive activity in the stallion, ram and hamster. Animal reproduction science 58:197–213.

Golder SA, Macy MW (2011) Diurnal and seasonal mood vary with work, sleep, and daylength across diverse cultures. Science 333:1878–1881.

Harmatz MG, Well AD, Overtree CE, Kawamura KY, Rosal M, Ockene IS (2000) Seasonal variation of depression and other moods: a longitudinal approach. J Biol Rhythms 15:344–350.

Hirvonen J, Aalto S, Lumme V, Någren K, Kajander J, Vilkman H, Hagelberg N, Oikonen V, Hietala J (2003) Measurement of striatal and thalamic dopamine D2 receptor binding with 11C-raclopride. Nuclear medicine communications 24:1207–1214.

Innis RBaCVJaDJaFMaGAaGRNaHJaHSaHSC (2007) Consensus nomenclature for in vivo imaging of reversibly binding radioligands. In, pp 1533–1539, pmid = 17519979.

Jameson AN, Siemann JK, Melchior J, Calipari ES, McMahon DG, Grueter BA (2023) Photoperiod Impacts Nucleus Accumbens Dopamine Dynamics. Eneuro 10.

Kaasinen V, Jokinen P, Joutsa J, Eskola O, Rinne JO (2012) Seasonality of striatal dopamine synthesis capacity in Parkinson’s disease. Neurosci Lett 530:80–84.

Karjalainen T, Tuisku J, Santavirta S, Kantonen T, Bucci M, Tuominen L, Hirvonen J, Hietala J, Rinne JO, Nummenmaa L (2020) Magia: Robust Automated Image Processing and Kinetic Modeling Toolbox for PET Neuroinformatics. Front Neuroinform 14:3.

Koskela A, Kauppinen T, Keski-Rahkonen A, Sihvola E, Kaprio J, Rissanen A, Ahonen A (2008) Brain Serotonin Transporter Binding of [123I] ADAM: Within-Subject Variation between Summer and Winter Data. Chronobiology international 25:657–665.

Lam RWaLRD (2000) Pathophysiology of seasonal affective disorder: A review. In: Journal of Psychiatry and Neuroscience, pp 469–480, pmid = 11109298.

Lammertsma AA, Hume SP (1996) Simplified reference tissue model for PET receptor studies. NeuroImage 4:153–158.

Levitan RD (2022) The chronobiology and neurobiology of winter seasonal affective disorder. Dialogues in clinical neuroscience.

Liu C, Kaeser PS (2019) Mechanisms and regulation of dopamine release. Current opinion in neurobiology 57:46–53.

Malén T, Karjalainen T, Isojärvi J, Vehtari A, Bürkner P-C, Putkinen V, Kaasinen V, Hietala J, Nuutila P, Rinne J (2022) Atlas of type 2 dopamine receptors in the human brain: Age and sex dependent variability in a large PET cohort. NeuroImage 255:119149.

Matheson GJ, Schain M, Almeida R, Lundberg J, Cselenyi Z, Borg J, Varrone A, Farde L, Cervenka S (2015) Diurnal and seasonal variation of the brain serotonin system in healthy male subjects. NeuroImage 112:225–231.

Meyer C, Muto V, Jaspar M, Kusse C, Lambot E, Chellappa SL, Degueldre C, Balteau E, Luxen A, Middleton B, Archer SN, Collette F, Dijk DJ, Phillips C, Maquet P, Vandewalle G (2016) Seasonality in human cognitive brain responses. Proceedings of the National Academy of Sciences of the United States of America 113:3066–3071.

Mooldijk SS, Licher S, Vernooij MW, Ikram MK, Ikram MA (2022) Seasonality of cognitive function in the general population: the Rotterdam Study. GeroScience:1–11.

Neumeister A, Willeit M, Praschak-Rieder N, Asenbaum S, Stastny J, Hilger E, Pirker W, Konstantinidis A, Kasper S (2001) Dopamine transporter availability in symptomatic depressed patients with seasonal affective disorder and healthy controls. Psychological medicine 31:1467–1473.

Pothos EN, Przedborski S, Davila V, Schmitz Y, Sulzer D (1998) D2-Like dopamine autoreceptor activation reduces quantal size in PC12 cells. Journal of Neuroscience 18:5575–5585.

Praschak-Rieder N, Willeit M, Wilson AA, Houle S, Meyer JH (2008) Seasonal variation in human brain serotonin transporter binding. Archives of general psychiatry 65:1072–1078.

Rolls ET, Joliot M, Tzourio-Mazoyer N (2015) Implementation of a new parcellation of the orbitofrontal cortex in the automated anatomical labeling atlas. NeuroImage 122:1–5.

Schmitz Y, Schmauss C, Sulzer D (2002) Altered dopamine release and uptake kinetics in mice lacking D2 receptors. Journal of Neuroscience 22:8002–8009.

Shahar DR, Froom P, Harari G, Yerushalmi N, Lubin F, Kristal-Boneh E (1999) Changes in dietary intake account for seasonal changes in cardiovascular disease risk factors. European journal of clinical nutrition 53:395–400.

Small DM, Jones-Gotman M, Dagher A (2003) Feeding-induced dopamine release in dorsal striatum correlates with meal pleasantness ratings in healthy human volunteers. NeuroImage 19:1709–1715.

Sowell ER, Peterson BS, Thompson PM, Welcome SE, Henkenius AL, Toga AW (2003) Mapping cortical change across the human life span. Nature neuroscience 6:309–315.

Spies M et al. (2018) Brain monoamine oxidase A in seasonal affective disorder and treatment with bright light therapy. Transl Psychiatry 8:198.

Spindelegger C, Stein P, Wadsak W, Fink M, Mitterhauser M, Moser U, Savli M, Mien L-K, Akimova E, Hahn A (2012) Light-dependent alteration of serotonin-1A receptor binding in cortical and subcortical limbic regions in the human brain. The World Journal of Biological Psychiatry 13:413–422.

Sulzer D, Cragg SJ, Rice ME (2016) Striatal dopamine neurotransmission: regulation of release and uptake. Basal ganglia 6:123–148.

Sun L, Laurila S, Lahesmaa M, Rebelos E, Virtanen KA, Schnabl K, Klingenspor M, Nummenmaa L, Nuutila P (2023) Secretin modulates appetite via brown adipose tissue-brain axis. Eur J Nucl Med Mol Imaging 50:1597–1606.

Sun L, Aarnio R, Herre EA, Karna S, Palani S, Virtanen H, Liljenback H, Virta J, Honkaniemi A, Oikonen V, Han C, Laurila S, Bucci M, Helin S, Yatkin E, Nummenmaa L, Nuutila P, Tang J, Roivainen A (2022) [(11)C]carfentanil PET imaging for studying the peripheral opioid system in vivo: effect of photoperiod on mu-opioid receptor availability in brown adipose tissue. Eur J Nucl Med Mol Imaging.

Sun L et al. (2021) Seasonal Variation in the Brain mu-Opioid Receptor Availability. The Journal of neuroscience : the official journal of the Society for Neuroscience 41:1265–1273.

Tsai H-Y, Chen KC, Yang YK, Chen PS, Yeh TL, Chiu NT, Lee IH (2011) Sunshine-exposure variation of human striatal dopamine D2/D3 receptor availability in healthy volunteers. Progress in Neuro-Psychopharmacology and Biological Psychiatry 35:107–110.

Wehr TA (1991) The durations of human melatonin secretion and sleep respond to changes in daylength (photoperiod). J Clin Endocrinol Metab 73:1276–1280.

Wu Q, Reith ME, Walker QD, Kuhn CM, Carroll FI, Garris PA (2002) Concurrent autoreceptor-mediated control of dopamine release and uptake during neurotransmission: an in vivo voltammetric study. Journal of Neuroscience 22:6272–6281.

Zhang R, Volkow ND (2023) Seasonality of brain function: role in psychiatric disorders. Translational Psychiatry 13:65.

Zisapel N (2001) Melatonin–dopamine interactions: from basic neurochemistry to a clinical setting. Cellular and molecular neurobiology 21:605–616.

